# Topographic complexity shapes adaptive genomic variation in co-occurring plant species

**DOI:** 10.64898/2026.08.07.743600

**Authors:** Simón E. Lobos, Collin W. Ahrens, Paul D. Rymer, Kathryn A. Hodgins, Adam D. Miller

## Abstract

Co-occurring species often face similar selective environments, although adaptive responses to these environments are generally assumed to be species-specific, particularly in complex landscapes where selective pressures are likely to be multi-dimensional. We test this assumption by contrasting genotype-environment associations (GEAs) among a range of co-occurring but unrelated plants with different life histories from an isolated, mountainous national park in south-eastern Australia. Analyses were performed using single nucleotide polymorphism (SNP) loci derived from reduced genome representation sequencing to investigate genomic associations with spatial and environmental drivers across unrelated plant species within the same heterogeneous landscape. Several species showed GEAs that aligned with similar environmental gradients, particularly edaphic features, suggesting that similar selective pressures can shape genomic responses across taxa. Other species exhibited distinct spatial and environmental associations, highlighting idiosyncratic outcomes. Notably, GEAs were detected at fine spatial scales despite generally low levels of genome-wide divergence, suggesting adaptive variants can persist in the face of gene flow under strong selective pressure. This study highlights how community-level genomic diversity is shaped by common environmental processes, with implications for biodiversity management in rapidly changing environments, where diverse ecosystem-level responses to selection may underpin resilience.

## 1. Introduction

The Earth’s biota is facing the greatest mass extinction episode in 65 million years (Cowie et al., 2022; Ceballos & Ehrlich, 2023), with around half of species currently undergoing strong range contractions (Finn et al., 2023). Addressing this global biodiversity crisis requires improved conservation policies, including genetic management plans, to quantify and predict evolutionary responses to anthropogenic climate change (Hoffmann & Sgrò, 2011; Harris et al., 2018; Flanagan et al., 2018). Empirical studies suggest that the resilience and adaptive potential of a species to novel threats are largely driven by pre-existing genetic variation (i.e., standing variation) for environment-relevant traits (Jump et al., 2009; Bragg et al., 2015; Catullo et al., 2019; Kardos et al., 2021). Conservation efforts that prioritise maintaining and enhancing genetic variation often focus on one or a few species due to practical limitations (Kardos et al., 2021; Zbinden et al., 2023). However, adopting a multi-species approach could uncover landscape-scale environmental features that structure communities within vulnerable ecosystems (Barrows et al., 2005; Schwenk & Donovan, 2011; Hand et al., 2015). Such insights could inform more comprehensive and systematic management strategies to better achieve long-term conservation goals, ultimately benefiting entire communities (Nielsen et al., 2017, 2020).

Species experiencing similar selective environments may exhibit distinct adaptive genetic responses to climate, due to differences in factors such as functional traits and ecological niche (Davis et al., 2005; Aitken et al., 2008; Tiffin & Ross-Ibarra, 2014). Variation in dispersal ability (Lenormand, 2002), mating system (Hodgins & Yeaman, 2019), generation time (Aitken et al., 2008), and phenotypic plasticity (Palacio-López et al., 2015; Levis & Pfennig, 2016; Blackman et al., 2017) can influence how species experience and respond to spatially varying selection. Species may also differ in which climatic variables exert the strongest selective pressures, depending on their functional traits, physiological tolerances, microhabitat use, and biotic interactions (Vargas et al., 2017; Chaturvedi et al., 2022; Chen et al., 2023; Voolstra et al., 2023). In complex environments where multiple abiotic and biotic factors can vary simultaneously, selection may act on different combinations of traits across space and time, depending on how species’ physiological traits and ecological strategies (e.g., microhabitat use, resource acquisition) mediate their responses to local conditions (Anderson et al., 2014; White et al., 2022). As a result, adaptive responses may diverge among species due to trait-specific selection regimes and potential trade-offs among traits that constrain multivariate adaptation (Schluter, 1996; Etterson & Shaw, 2001). Such trade-offs often arise from pleiotropic constraints, in which genetically correlated traits cannot evolve independently, thereby shaping the direction and pace of adaptation (Wagner & Zhang, 2011; Battlay et al., 2024).

The genetic architecture of adaptation may vary among species, with some exhibiting polygenic responses involving many small-effect loci, while others are shaped by fewer loci of larger effect (Yeaman, 2015; Boyle et al., 2017; Connallon & Hodgins, 2021). Differences in genome structure, such as recombination landscapes or structural variants, may further shape these patterns by influencing how adaptive loci are distributed across the genome and how strongly they are linked (Yeaman, 2013; Wellenreuther & Bernatchez, 2018). Finally, demographic history, gene flow, levels of standing genetic variation, and the underlying genetic architecture of climate-relevant traits can all influence the biological signal and statistical detectability of climate-associated loci (Lotterhos & Whitlock, 2015; Hoban et al., 2016; Hodgins & Yeaman, 2019), contributing to species-specific patterns even under shared environmental conditions.

Despite these complexities, evidence suggests that environmental gradients frequently shape adaptive differentiation in co-occurring taxa (Raeymaekers et al., 2017; Walters et al., 2021; Yadav et al., 2021) and even lead to convergence in the genetic basis of adaptation among some species (Whiting et al., 2024). Consequently, even distantly related taxa can show broadly repeatable adaptive responses to similar selective environments (Ahrens et al., 2025). Even so, landscape genomic studies are needed to determine whether adaptive evolution is consistent and predictable when independent lineages in the same habitat experience similar landscape and environmental processes. Such multi-species assessments are crucial in conservation planning because they can inform strategies that prioritise the maintenance or enhancement of genetic variation across communities, ensuring sympatric species can respond to novel environmental stresses.

A landscape genomics framework offers a powerful approach to assess the molecular basis of local adaptation and identify selective drivers in non-model species (e.g., Bohutínská et al., 2021; Chaturvedi et al., 2022; Yeaman et al., 2016). Landscape genomics integrates population genomics, landscape ecology, and spatial statistics to reveal genome-wide patterns of neutral and adaptive variation shaped by environmental heterogeneity (Sork et al., 2013; Shryock et al., 2015; Balkenhol et al., 2017; Jia et al., 2020). It may additionally be applied to predict adaptive capacity by quantifying gene flow among locally adapted populations distributed across environmental gradients (Aitken & Whitlock, 2013; Sork, 2016). From the perspective of conservation biology, these approaches are important for detecting adaptation to local environmental conditions, delineating adaptive units, and mitigating risks of maladaptation (Savolainen et al., 2007; Carvalho et al., 2021). Plants are particularly well-suited to landscape genomic analyses due to their sessile nature, high levels of genomic and phenotypic diversity, and variable dispersal strategies (Corlett & Westcott, 2013). Furthermore, many plant species, particularly long-lived taxa, have limited capacity to track rapidly shifting climates, underscoring the importance of landscape genomics research for understanding their evolutionary potential (Forester et al., 2016; Oldfather et al., 2020).

In this study, we identified putative climate-driven adaptive genomic variation in six widespread, co-occurring, but taxonomically unrelated plant species across the heterogeneous landscape of Gariwerd-Grampians National Park (hereafter “Gariwerd”), south-eastern Australia. This is a unique study that sets out to determine whether unrelated species with different life histories, co-occurring in complex landscapes, show genomic convergence in the face of common selective environments. We take advantage of the fact that complex landscapes can decouple patterns of neutral and adaptive variation, enhancing the detection of environment-associated loci (Lotterhos & Whitlock, 2015). Specifically, we used reduced-representation sequencing to conduct genotype-environment association (GEA) analyses aimed at addressing two questions: (i) Do multiple co-occurring species show genomic signals of adaptation to local environments? (ii) Are the spatial patterns and environmental drivers of putative adaptive genomic variation consistent across taxa? We examine how both shared and contrasting patterns of adaptive variation contribute to community resilience and ecosystem function, and discuss their implications for biodiversity management under rapid environmental change. We further integrate these insights with known patterns of gene flow to help prioritise actions that maximise community-wide evolutionary resilience.

## 2. Materials and Methods

### 2.1 Study site

The Gariwerd mountains (First Nations Jardwadjali name for Grampians National Park; Wilkie, 2020) are situated in the central western part of Victoria, Australia (Figure 1). They comprise a series of rugged, sub-parallel Devonian sandstone ranges, separated by alluvial valleys and sand sheets (Pollock et al., 2013). The region experiences a semi-Mediterranean climate, with hot and dry summers and mild, rainy winters (Enright et al., 1994). Environmental variation across the landscape is strongly structured along elevational gradients, which integrates sharp variation in temperature, precipitation, and edaphic features (Table S1). Such heterogeneity has been shown to strongly influence plant community composition, especially in relation to soil nutrients, water availability, aspect, and fire regime within the region (Enright et al., 1994; Pollock et al., 2015).

**Figure 1.**
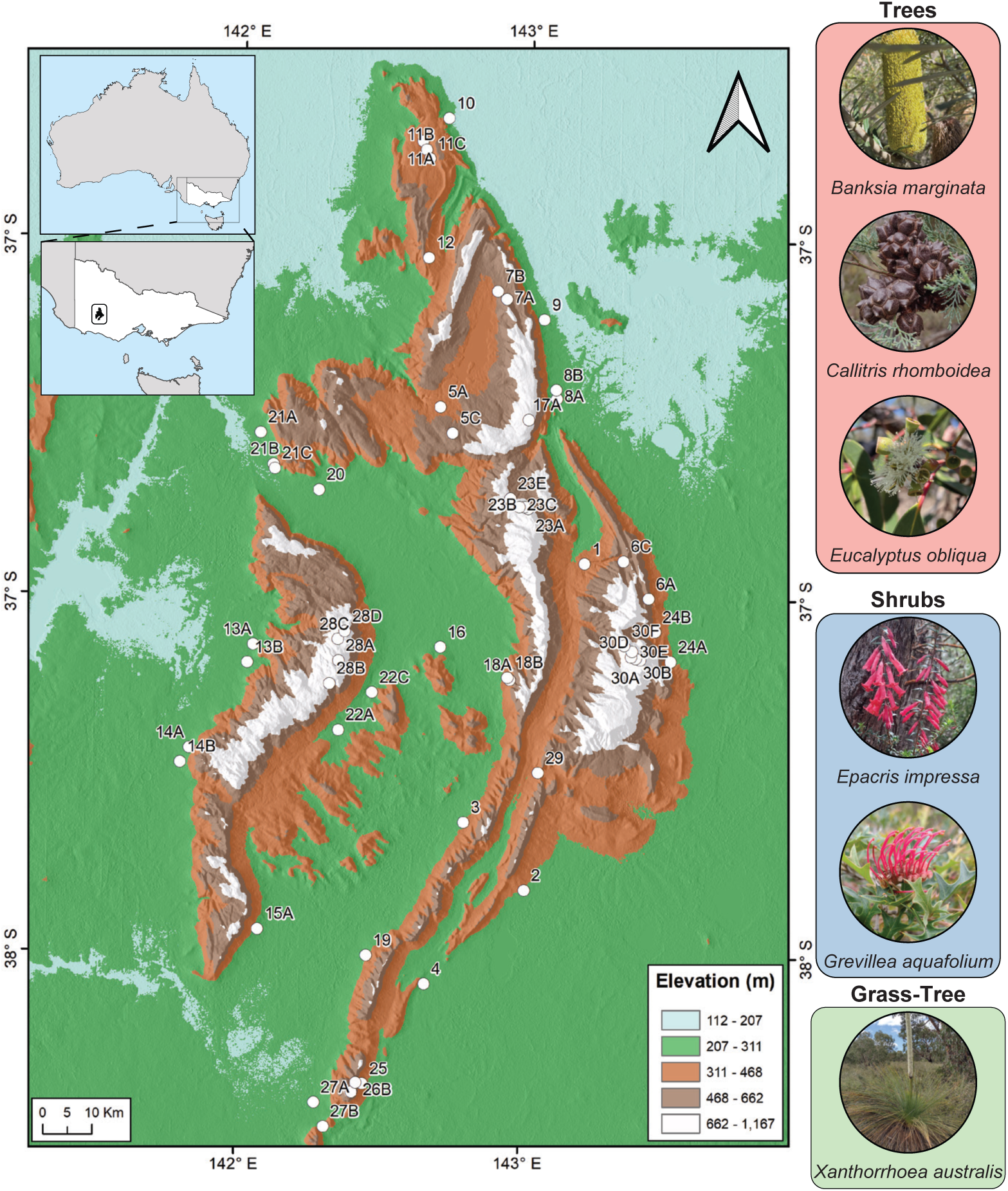
Map of sampling locations used for estimating patterns of genetic-environment associations among flora populations spanning Gariwerd.

### 2.2 Species and site selection

Six widespread, co-occurring plant species were selected for comparative landscape genomic analysis. These consisted of three trees (*Banksia marginata* Cav., *Callitris rhomboidea* R.Br. ex Rich., *Eucalyptus obliqua* L’Hér.), two shrubs (*Epacris impressa* Labill., *Grevillea aquifolium* Lindl.), and a single grass-tree species (*Xanthorrhoea australis* R.Br.; Table S1). For simplicity, species will be hereafter referred to by their genus only. These species represent a variety of life-forms, dispersal and pollination strategies (Table S1) that capture a broad representation of the region’s common flora (Costermans, 2007). Each species was expected to be diploid based on inferences from species-specific karyotypes or those from related taxa (Rice et al., 2015).

We adopted a hierarchical, replicated sampling regime to capture locations representing both extreme and intermediate habitats across key environmental gradients, particularly temperature, precipitation and soil properties (Table S2; Figure S1). This design also ensured that habitat types were replicated across different areas of Gariwerd, to help control for genetic variation shaped solely by demography (e.g., gene flow and drift). A total of 55 sampling locations across Gariwerd were selected, allowing for sampling of each species from an average of 29 sampling locations based on species presence/absence (Figure 1; Table S2).

### 2.3 Sample collection

At each site, 3–5 leaves were collected from 10–15 individuals per species between October 2021 and December 2022 (Figure 1; Table S2). Leaf tissue was preserved on coffee filter paper and desiccated with silica gel to minimise DNA degradation. Geographic coordinates were recorded at each location using a Garmin GPSMAP 78 handheld GPS device (Garmin Ltd., Schaffhausen, Switzerland). Genetic samples were stored at room temperature prior to genomic processing. We then weighed leaf samples (∼10–15 mg) from six individuals per species in each population, totalling 1,092 individuals (Table S2).

### 2.4 DNA isolation, library preparation and sequencing

Silica-dried leaf samples were sent to Diversity Arrays Technology (DArT Pty Ltd, Canberra, Australia) for DNA extraction, library preparation, sequencing and SNP genotyping using proprietary DArTseq^TM^ technology. DArTseq^TM^ is similar to the Restricted-site Associated DNA sequencing (RAD-seq) method and has been widely utilised to generate single nucleotide polymorphism (SNP) datasets across diverse plant species (Steane et al., 2015; Jordan et al., 2017; Rutherford et al., 2018; Bradbury et al., 2019; Robins et al., 2021, 2023; Wilson et al., 2022). This approach enables targeted selection of genome fractions enriched for active genes, which may be associated with diverse traits of ecological and evolutionary relevance in plants (Nadeem et al., 2018). Data quality is enhanced by sequencing multiple libraries per sample, allowing the calculation of reproducibility scores for each candidate marker and ensuring reduced missing data and more reliable SNP calls (Rutherford et al., 2021).

The DArTseq protocol has been described in detail by Sansaloni et al. (2011) and Kilian et al. (2012). Briefly, DNA was extracted by DArT using a NucleoMag 96 Tissue Kit (Macherey-Nagel, Germany) coupled with NucleoMag SEP (Ref. 744900) to allow automated separation of high-quality DNA on a Tecan Freedom EVO liquid handler (Männedorf, Switzerland) (Georges et al., 2018). Reduced representation libraries were generated using restriction enzyme combinations tailored to each taxon: *PstI* + *HpalI* for *Banksia* and *Eucalyptus*, and *PstI* and *MseI* for *Callitris*, *Epacris*, *Grevillea* and *Xanthorrhoea*. Specialised adaptors were then ligated to the digested DNA, and ligated fragments were amplified by PCR (1 min 94 °C; 30 cycles: 20 s 94 °C, 30 s 58 °C, 45 s 72 °C; with 7 min extension at 72 °C). After PCR, the amplified products were standardised and pooled for sequencing on an Illumina HiSeq2500. Raw sequence data were processed using a proprietary DArT analytical pipeline (*DArTsoft14*), which filtered poor-quality sequences, SNP calling and generated final genotypes. To allow a measure of technical repeatability in genotype calls and library preparation, 30% of samples were sequenced again to ensure the quality of the data. Additionally, the technical replicates were used to model the Mendelian distribution of alleles to discriminate true allelic variants from paralogous regions and remove sequencing errors (Sansaloni et al., 2011; Kilian et al., 2012).

### 2.5 SNP quality control filtering

Further filtering of SNP loci was performed separately for each species using the *dartR* v.2.7.2 package (Mijangos et al., 2022) in R version 4.4.0 (R Core Team, 2024). Potential clone mates were identified by assessing the proportion of shared alleles between pairs of individuals within each species dataset using the ‘gl.propShared’ function in *dartR*. Genetic thresholds for assignment were determined using pairwise shared-allele frequencies between accessions with a sequenced technical replicate. A threshold of 95–99% genetic similarity was used to assign individuals to clone groups, but assignments were adjusted following visual inspection of a neighbour-joining tree generated in the R package *ape* v.5.8 (Paradis & Schliep, 2019). Only one representative of each clonal group was retained for downstream filtering. Following Ahrens et al. (2021), we employed relaxed thresholds for missing data and minor allele frequency to increase the proportion of the genome interrogated for signatures of selection, since only a small fraction is expected to contribute to adaptive divergence. SNPs were filtered as follows: minor allele frequency cut-off of 2%, SNPs with a call rate of non-missing values below 80% for loci and individuals, and similar loci were removed due to the presence of paralogous sequences (hamming distance < 0.2). To minimise the effects of physical linkage, we retained only one SNP per contig, identified using its unique CloneID (i.e., the unique sequence tag identifier for each locus). We used the same parameters for each species to facilitate comparison.

### 2.6 Environmental data collection

Environmental data for each sampling location were retrieved from open-source GIS databases in raster format to represent bioclimatic, edaphic, and hydrological gradients (Figure S1). The environmental characteristics of each sampling location were quantified using a set of six relevant predictors, based on past work defining them as eco-physiologically important (Williams et al., 2012; Guerin et al., 2022; Mokany et al., 2022; Westerband et al., 2023), and common drivers of adaptive genomic variation in plant species (Yoder et al., 2014; Ahrens et al., 2018, 2019, 2021; Mostert-O’Neill et al., 2021; Schmidt et al., 2021; Filipe et al., 2022; Oyanoghafo et al., 2023). Bioclimatic information was obtained from the WorldClim v.2.06 database (Fick & Hijmans, 2017) as gridded files at 30 arc-second resolution (∼ 1 km^2^) for the years 1970–2000. Specifically, we used four bioclimatic variables, including annual mean temperature (*T*_MA_), maximum temperature of the warmest month (*T*_MAX_), annual precipitation (*P*_MA_) and precipitation of the driest month (*P*_MIN_). Soil variables were obtained from the TERN Soil and Landscape Grid of Australia database (https://data.csiro.au/; Grundy et al., 2015). We used two soil variables (available phosphorus–*S*_AP_ and soil pH in water and CaCl_2_–*S*_pH_) at 3 arc-second resolution (∼ 90 x 90 m pixels) and 0–5cm soil depth, as plant roots are mainly distributed in surface soils and because there are very strong correlations across depths (W. Luo et al., 2021). Finally, to take into account the hydrological characteristics of the landscape, we used the topographic wetness index (*TWI*), considered as a proxy for soil moisture conditions. *TWI* represents local topographic influence on hydrological processes based on catchment area and local slope inclination (Kopecký et al., 2021). *TWI* derived at 10m resolution from digital elevation models (DEMs) was downloaded through the CSIRO Data Access Portal (https://data.csiro.au/collections; Gallant & Austin, 2015). To account for the differing spatial resolutions, the rasters were calibrated and downscaled to the same resolution at a grid cell size of 30″ × 30″ using the packages *raster* v.3.6-26 (Hijmans, 2023) and *rgdal* v.1.6-7 in R (Bivand et al., 2006-2023).

Collinearity among all predictor variables was estimated by applying a variance inflation factor (VIF) test (Marquardt, 1970) implemented in the ‘vifcor’ function in the R package *usdm* v.2.1-7 (Naimi et al., 2014). This approach quantifies VIF for all variables and identifies variables with the highest correlation value. A correlation coefficient threshold of 0.6 was used following the recommendations of Sales et al. (2021) recommendations. Strong collinearity was observed among all bioclimatic predictor variables, so only *T*_MAX_ was retained for subsequent analyses. In contrast, all soil and hydrology variables were retained as none exhibited strong collinearity.

### 2.7 Genotype-environment associations

To identify putative genomic signatures of selection, we performed genotype-environment association analyses (GEAs) by correlating population-allele frequencies with location-specific environmental variables (Rellstab et al., 2015; Hoban et al., 2016; Ahrens et al., 2018). GEAs have been shown to outperform other analytical frameworks in recovering genomic regions under selection by environmental gradients (Rellstab et al., 2015). Analyses were executed separately for each species by employing two complementary individual-based GEA approaches: (i) LFMM2 (Caye et al., 2019) and (ii) BayPass v2.3 (Gautier, 2015). These univariate methods test one locus and one predictor variable at a time while, in different ways, overcoming the confounding effects of population structure (Feng & Du, 2022). LFMM2 estimates GEAs when simultaneously correcting for population structure with latent factors, while BayPass uses a neutral covariance matrix constructed from population allele frequencies. Therefore, a comprehensive adaptation test can be conducted using these two flexible and powerful approaches given an underlying pattern of population structure and environmental variation (Gautier, 2015; Ahrens et al., 2021; Alshwairikh et al., 2021; Filipe et al., 2022). For a more extensive discussion and details, see Ahrens et al. (2018, 2021), Lotterhos (2019, 2023) and L. Luo et al. (2021).

We first used latent factor mixed models (LFMM) to test for linear relationships between individual-based allele frequencies and environmental variables with random latent factors using a least-square method (Caye et al., 2019). We explored population structure by inferring individual ancestry coefficients representing the proportions of each individual’s genome that originated from multiple ancestral gene pools. Calculations were performed with the sparse non-negative matrix factorisation (sNMF) method implemented in the ‘snmf’ function in the R package *LEA* v.3.10.2 (Frichot & François, 2015). The sNMF method is comparable to ADMIXTURE and STRUCTURE programs but outperforms them in estimating homozygote and heterozygote frequencies and does not rely on Hardy-Weinberg equilibrium assumptions (Frichot et al., 2014). Ancestry coefficient values were defined between 1 (i.e., a single panmictic population) and 30 ancestral populations (*K*) by generating an entropy criterion that evaluates the fit of the statistical model to the data using a cross-validation technique (Frichot & François, 2015). We selected the *K* value with the lowest cross-entropy value using 100 repetitions and 100 as the alpha regularisation parameter. Subsequently, the optimal *K* factor for each taxon (Figure S2) was used to inform the LFMM2 to identify whether allele frequencies were correlated with any of the environmental variables. To increase the statistical power of associations and adhere to LFMM2’s requirement of a complete dataset, missing genotype data were imputed via the ‘impute’ function of the *LEA* package, using the most common allele frequency observed in each *K* with the method ‘mode’. Next, we used the function ‘lfmm_ridge’ to compute a regularised least squares estimate using a ridge penalty. Individual associations between each SNP frequency and each environmental variable were assessed using a statistical test calibrated using the genomic inflation factor (function ‘lfmm_test’). This function performs associations between environmental variables on each SNP, and *p*-values are calibrated with the genomic inflation factor method. Corrections for multiple comparisons were applied with the Benjamin-Hochberg algorithm with a false discovery rate (FDR) threshold of 5% (Benjamini & Hochberg, 1995). Significant associations were determined using an α threshold of 0.001 after applying a false discovery rate (FDR) adjustment for 5% (Ahrens et al., 2021).

Second, we explored GEAs with BayPass, which identifies environment-driven differences in allele frequencies between populations. BayPass is based on an improved version of the Bayesian hierarchical model initially proposed in BayEnv (Coop et al., 2010; Gautier, 2015), and consists of a two-step process. First, the core model (without the environmental data) was run to estimate a covariance matrix (Ω) of population allele frequencies, which is an approximation of genomic differentiation between populations caused by shared evolutionary history (Gautier, 2015). To reach convergence and reproducibility of the MCMC estimates, five independent runs, each with a randomly chosen seed, were performed using default parameters, except for the number of sampled parameter values, which was set to 5,000, and the burn-in period length, which was set to 50,000 iterations. Second, to detect evidence of genotype-environment correlations corrected for population structure, the auxiliary model was run using the average of the five covariance matrices as input and the four environmental variables. Environmental variables were scaled using the “-scalecov” option, and the same running parameters as the core model were applied. The strength of the association between genotype and the covariates was assessed by calculating the average of the log-transformed Bayes Factor (BF) in deciban units (dB) for each locus and environmental predictor. Significance was determined following Jeffrey’s criterion for decisive evidence (BF ≥ 20; Jeffreys, 1961). Finally, UpSet plots were generated to depict the results of the GEAs, showing the strength of associations between individual loci and environmental variables per species. The plots were generated with the Customised UpSet Plots codes provided by Chenxin Li [https://github.com/cxli233/customized_upset_plots].

### 2.8 Predicting spatial patterns and environmental drivers of adaptive genomic variation

Gradient Forest (GF) was used to predict areas of high adaptive genetic variation and important environmental drivers within Gariwerd. The GF model was originally developed for community ecology applications and specifically to detect species turnover patterns in response to environmental gradients (Ellis et al., 2012). GF has been adapted to a landscape genomics framework to associate genomic variation with current and future environmental gradients (Fitzpatrick & Keller, 2015). GF is a flexible modelling approach implemented in the R package *gradientForest* v.0.1-37 (Ellis et al., 2012), that uses a regression tree-based machine learning algorithm to fit non-linear relationships between genomic data to environmental gradients (Capblancq et al., 2020). Furthermore, the algorithm recursively partitions allele frequencies at numerous split values along each environmental gradient and calculates the change in allele frequencies for each split. The split importance (i.e., the amount of genomic variation explained by each split value) is cumulatively summed along the environmental gradient and aggregated across alleles to build a non-linear turnover function that identifies loci significantly influenced by the predictor variable (Ellis et al., 2012).

We performed GF model analyses on candidate adaptive loci from the GEAs of four environmental variables identified using LFMM2 and BayPass. Analyses were run over 500 regression trees for each environmental variable, with all other parameters set to default settings. The contribution of each candidate locus was weighted by the coefficient of determination (*R*^2^) of its environmental association. These weights were used to summarise turnover and estimate the relative importance of environmental predictors in explaining allele-frequency variation. We conducted a principal component analysis (PCA) using the R function ‘prcomp’ to summarise the GF-predicted multidimensional genomic variation for each species. The top three principal components (PCs) were assigned to a red-green-blue colour palette for the biplot (Fitzpatrick & Keller, 2015). In the biplot, similar colours in the sampled space correspond to similar allele frequencies. Finally, the spatial distribution of environmentally associated allelic variation for each species was mapped across the study area using the *ggplot2* v.4.0.1 R package (Wickham, 2011), where changes in background colour represent the spatial turnover of putatively adaptive alleles.

To quantify the concordance between gradient forest prediction spaces across species pairs, PCAs of predicted allele frequencies were superimposed by the Procrustes analysis using the function ‘procrustes’ in the *vegan* v.2.6-4 R package (Oksanen et al., 2018). This multivariate method rotates and scales the axes of the first three principal components in an Euclidean space while minimising the sum of squares differences between two or more datasets (Peres-Neto & Jackson, 2001). The absolute distance between locations in the genetic space and the rotated ordination space (i.e., Procrustes residuals) was then used as a measurement of prediction uncertainty. Then, the Procrustes residuals were mapped to visualise spatial differences in genetic composition across all pairwise comparisons of the six species using the *tmap* v.4.2 R package (Tennekes, 2018) (extracted to a 90-m^2^ point grid overlaid on the study region).

### 2.9 Inferring similarities in genetic variation patterns between species

To summarise similarities in environmental-genetic responses across species, we conducted a standard linear regression with Procrustes residuals, with predicted adaptive structure in space as the dependent variable and the pairwise correlation coefficients between the *R*^2^-weighted importance values of each environmental variable as the predictors, utilising the function ‘lm’ in the stats package. Pairwise Pearson’s correlation coefficients were calculated using the matrix of *R*^2^ per species and environmental variables on each species pair combination using the function ‘cor’ in R.

## 3. Results

High-density DArTseq genotyping yielded a mean of 106,382 biallelic SNPs per species (range 20,212–190,026; SD 60,397; Table S3). After quality filtering, a mean of 18,324 high-quality SNPs (range 2,521–35,082; SD 13,003) remained for population genomic analyses, representing an average of 153 individuals (range 132–174; SD 17) sampled across 29 locations (range 27–30; SD 1) per species (Table S3).

### 3.1 Genotype-environment associations

GEA identified 1,416 candidate SNPs across all species (∼236 per species), representing ∼1% of the total SNPs analysed (Table S4). Of these, 1,289 loci (91%) were uniquely associated with a single environmental predictor (Table S4). We defined candidates as loci found to be significant by either LFMM2 or BayPass. Concordance between methods was low, with <11% of candidates overlapping between LFMM2 and BayPass for each species (Table S4). Candidate counts were broadly similar across predictors, with the largest number associated with *S*_AP_ (413 loci; 29%) and the fewest with *S*_pH_ (312; 22%) (Table S4). LFMM2 yielded more candidates than BayPass overall (71% of all candidates). Among LFMM2 candidates, most were linked to *T*_MAX_ (31%) (Table S4). In contrast, BayPass candidates were enriched for soil and hydrology variables, primarily *S*_AP_ (35%) and *TWI* (31%).

Candidate counts varied among species (Table S4; Figures S3-S8). *Banksia* showed 30 candidate SNPs (1% of all SNPs), with most associated with *S*_pH_ (33%). *Callitris* yielded 158 candidates (1% of all SNPs), most of which were associated with *TWI* and *S*_AP_ (32% and 30%, respectively). *Epacris* displayed 281 candidates (2%), dominated by *S*_pH_ (33%) and *T*_MAX_ (31%). For *Eucalyptus*, the majority of the 116 candidates (1%) were linked to *T*_MAX_ (42%), whereas *Grevillea* had 600 candidates (2%), primarily explained by associations with *S*_AP_ (38%). Finally, *Xanthorrhoea* had 231 candidates (1%), with similar counts across environmental predictors; *S*_pH_ (28%), *T*_MAX_ and *S*_AP_ (26%), *TWI* (21%) (Table S4; Figures S3-S8).

### 3.2 Predicting spatial patterns and environmental drivers of adaptive genomic variation

Gradient Forest (GF) analyses retained 699 SNPs with positive predictive power (*R*² > 0) across the six species (Table S5), representing 54% of the 1,289 candidate loci detected by GEA analyses. The highest proportion of candidate SNPs with predictive power was observed in *Epacris* (mean *R*^2^ = 0.066; 73% of SNPs) and *Eucalyptus* (mean *R*^2^ = 0.063; 61% of SNPs). In contrast, the lowest proportion of candidate loci with predictive power was observed in *Callitris* (mean *R*^2^ = 0.051; 53% of SNPs) and *Grevillea* (mean *R*^2^ = 0.049; 45% of SNPs) (Table S5). *Xanthorrhoea* (mean *R*^2^ = 0.053; 58% of SNPs) and *Banksia* (mean *R*^2^ = 0.038; 57% of SNPs) had an intermediate but similar proportion of candidate loci with predictive power.

Allelic turnover at candidate loci was most strongly influenced by *TWI* in *Callitris*, *Eucalyptus* and *Xanthorrhoea*, *S*_AP_ in *Banksia* and *Grevillea* and *S*_pH_ in *Epacris* (Figure 2A). Overall, *Epacris* and *Eucalyptus* showed the strongest *R*²-weighted importance across predictors (except *S*_AP_), whereas *Banksia* exhibited generally weak associations across variables (except *S*_AP_) (Figure 2). These patterns were reflected in the GF cumulative importance curves (Figure 2B). *Epacris* and *Eucalyptus* reached the greatest maximum cumulative turnover with similarly shaped curves across predictors (except for *S*_AP_ in *Epacris*; Figure 2B), while *Callitris*, *Grevillea*, and *Xanthorrhoea* showed comparable turnover profiles for *S*_pH_ and *T*_MAX_ (Figure 2B). In contrast, *Banksia* displayed flatter curves indicative of lower turnover across all predictor variables (Figure 2B). Across all species, turnover in allele frequencies occurred more quickly in cooler areas, with more acidic soils, higher phosphorus levels, and lower topographic wetness (Figure 2B).

**Figure 2.**
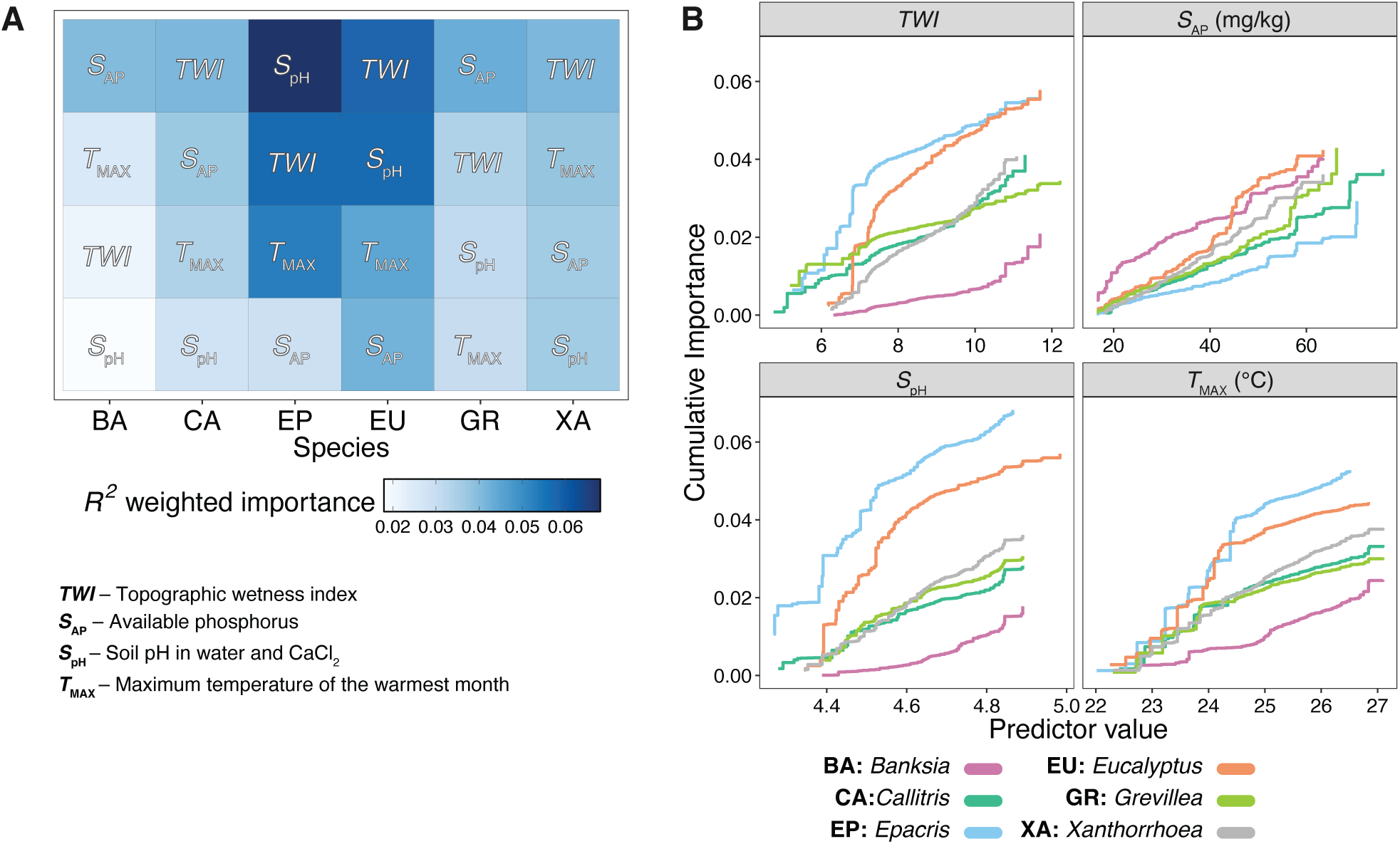
Outcomes of gradient forest models. (A) The relative importance of each environmental predictor variable in describing turnover in allele frequencies across candidate loci for each species. Darker blue squares indicate greater relative importance of predictor variables. All predictor variables were ranked by importance weighted by *R*^2^ for each species. (B) Cumulative importance curves showing the overall pattern of genomic compositional change (*R*^2^, y-axis) for each environmental gradient (x-axis). Turnover functions for each curve are aggregated across candidate loci. The maximum height of each curve indicates the relative amount of allelic turnover and the relative importance of each environmental variable.

Across all species, spatial turnover of putatively adaptive allele frequencies was steepest along elevation gradients, where habitat conditions (temperature, water availability, soil chemistry) shift abruptly over hundreds of metres (Figure 3). The strongest predicted turnover occurred among high-elevation sites near the centre of Gariwerd (i.e., Boroka Lookout– site 17A; Mt Rosea– site 23; Mt Thackeray– site 28 and Mt William– site 30; Table S2, Figure 3A– F), which is reflected by the clustering of locations in the principal components analysis of GF transformed environmental variables (Figure 3G–L). These locations are characterised by lower temperature (*T*_MAX_), lower water availability (*TWI*), high phosphorus (*S*_AP_), and lower soil pH (*S*_pH_). Nonetheless, subtle species-specific differences (Figure 3A–F) suggest that co-occurring taxa respond to the same landscape through partially distinct genomic responses. Each point in the PCA represents the principal component (PC) score for locations in the species distribution range.

**Figure 3.**
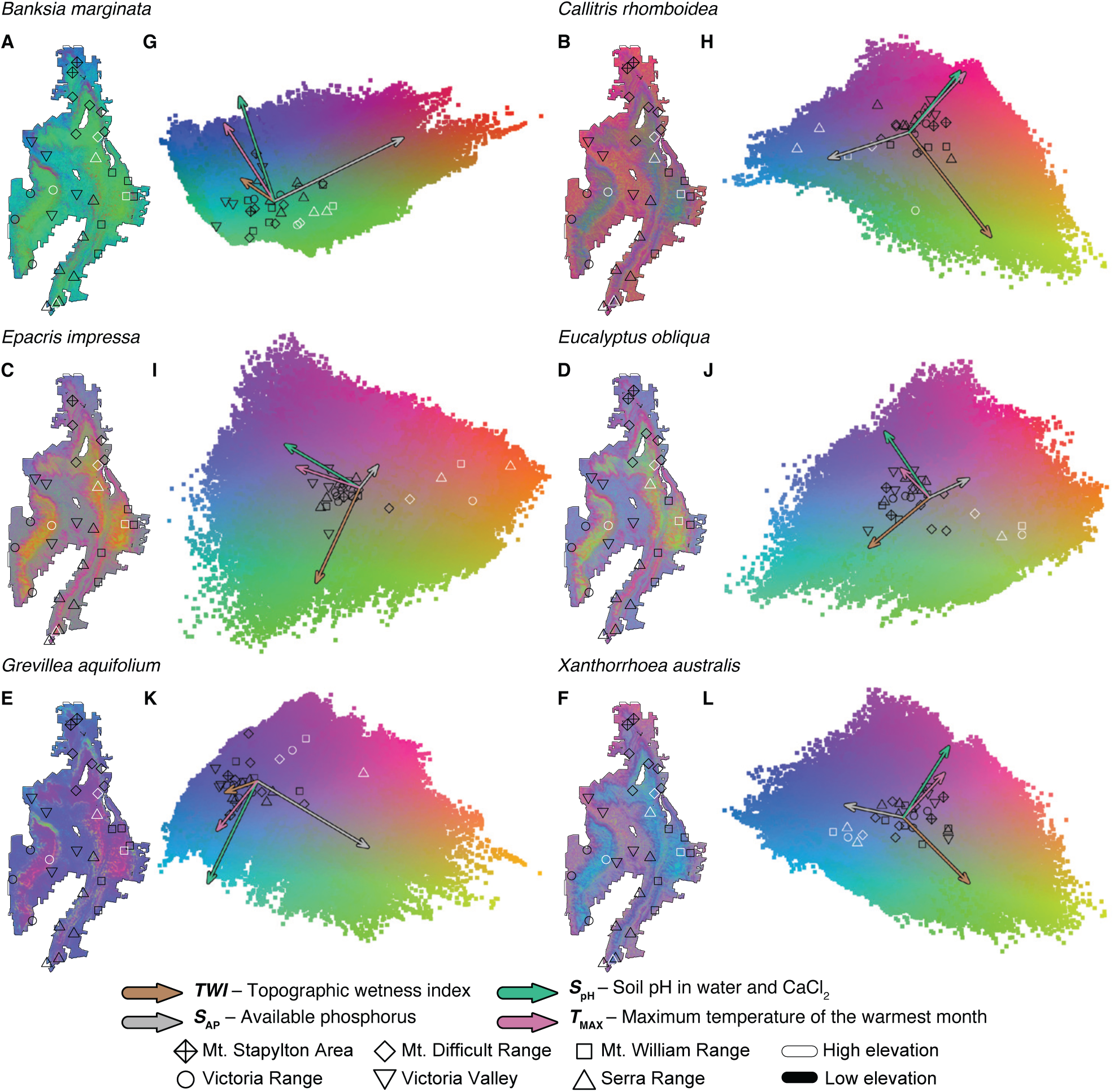
The composition of genomic turnover of each species from the Gradient Forest analysis for candidate loci. (A–F) Predicted spatial turnover in allele frequencies interpolated across the study area, with different symbols indicating sampling locations according to the area of origin. Regions with similar colours are expected to contain populations with comparable genetic composition, whereas regions with different colours suggest varying genotypes. (G–L) Biplots of the first two principal dimensions of the biologically transformed environment space illustrate the influence of environmental variation on allele frequencies. Background colours are based on the first three principal components of transformed environmental variables, with vector colour-coded indicating the direction and magnitude of major environmental variables.

Pairwise Procrustes residual analysis demonstrated diverse spatial patterns of allelic turnover between most species pairs, which were grouped into low (L), intermediate (I), and high (H) differentiation categories (Figure 4; Table S7). The lowest differentiation, based on Procrustes residual values, was observed between *Callitris* and *Xanthorrhoea* (*t_o_* = 0.009; Figure 4A), *Banksia* and *Grevillea* (*t_o_* = 0.017; Figure 4B, Table S7), *Eucalyptus* and *Xanthorrhoea* (*t_o_* = 0.018; Figure 4C, Table S7), *Callitris* and *Eucalyptus* (*t*_o_ = 0.021; Figure 4D, Table S7), and *Banksia* and *Callitris* (*t*_o_ = 0.025; Figure 4E, Table S7). These species (except *Eucalyptus*) also tended to show lower amounts of change in the cumulative importance for most of the predictor variables and tended to be more similar. Intermediate Procrustes residual values were found between *Banksia* and *Xanthorrhoea* (*t_o_* = 0.031; Figure 4F, Table S7), *Epacris* and *Xanthorrhoea* (*t_o_* = 0.032; Figure 4G, Table S7), *Epacris* and *Eucalyptus* (*t_o_* = 0.032; Figure 4H, Table S7), *Callitris* and *Epacris* (*t_o_* = 0.034; Figure 4I, Table S7), *Grevillea* and *Xanthorrhoea* (*t_o_* = 0.037; Figure 4J, Table S7), *Callitris* and *Grevillea* (*t_o_* = 0.038; Figure 4K, Table S7), and *Banksia* and *Eucalyptus* (*t_o_* = 0.044; Figure 4L, Table S7). Finally, the highest Procrustes residual values were obtained when comparing *Grevillea* with both *Eucalyptus* (*t_o_* = 0.061; Figure 4O, Table S7) and *Epacris* (*t_o_* = 0.059; Figure 4N, Table S7), followed by comparisons between *Banksia* and *Epacris* (*t_o_* = 0.054; Figure 4M, Table S7). These differences were notably strong in steep sloped areas across the main four ranges in the park (i.e., Mount Difficult Range, Serra Range, Victoria Range and Mount William Range).

**Figure 4.**
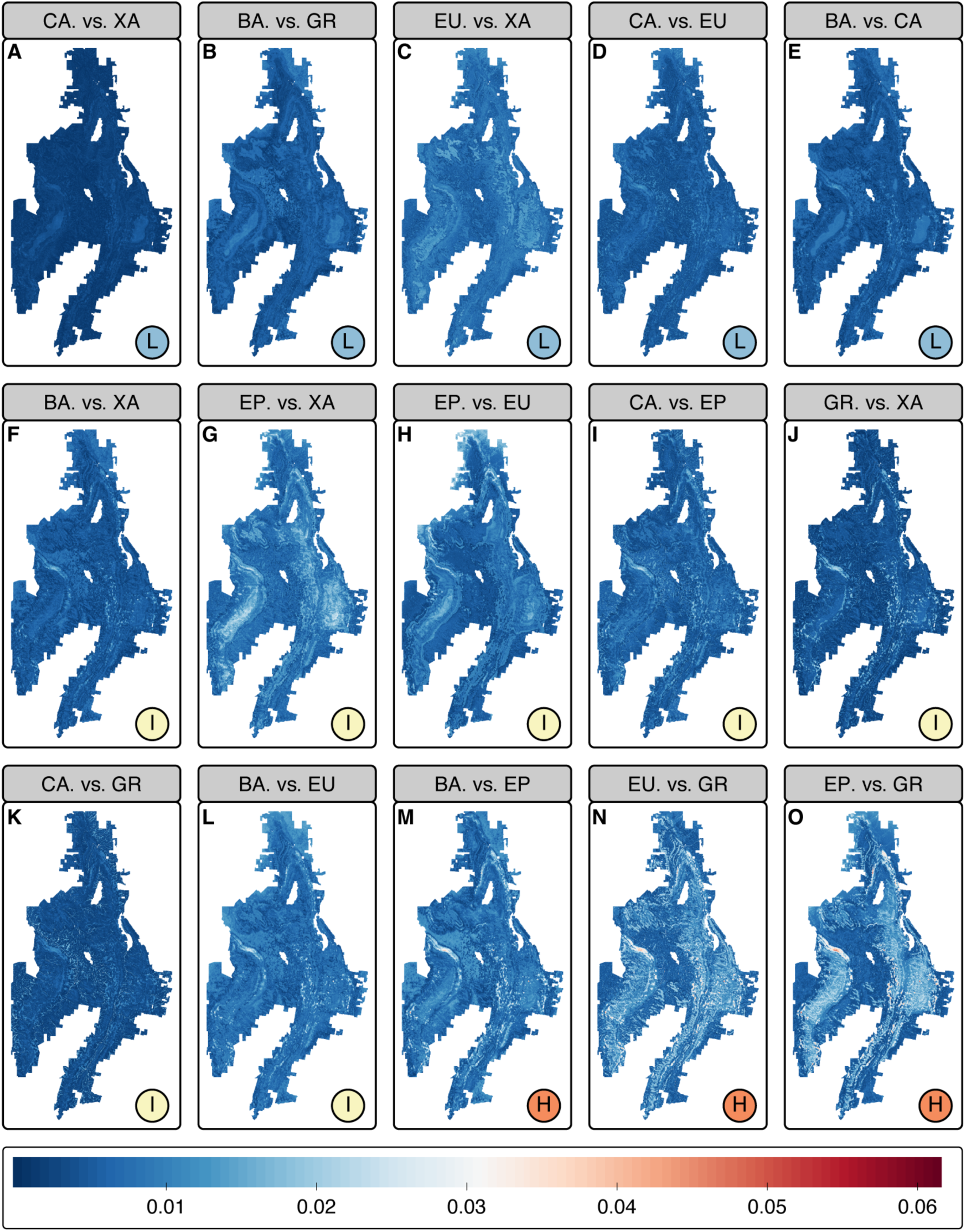
Pairwise comparisons of mapped predicted turnover in allele frequencies from gradient forest models based on Procrustes residuals (PR). Colours indicate areas with low (blue) to high (red) differences in predicted geographic patterns between species. Letters in circles represent the three main groups to describe concordance between gradient forest prediction spaces across species pairs (L–low, I–intermediate, and H–high differentiation). Labels indicate the corresponding species pairs. Species abbreviations are as follows: BA, *Banksia*; CA, *Callitris*; EP, *Epacris*; EU, *Eucalyptus*; GR, *Grevillea*; and XA, *Xanthorrhoea*.

### 3.3 Similarities in putative adaptive genetic variation patterns between species

A clear negative relationship was detected between similarity in environmental drivers of adaptive genetic variation (*R*^2^) and differences in predicted spatial patterns of genetic structure (PR) across species pairs (*p* = 0.001, *R*^2^ = 0.58; Figure 5). Species pairs with more similar environmental responses (high correlation in R²) tended to have similar spatial patterns of genetic structure (low PR) (e.g., *Banksia* vs *Grevillea* and *Callitris* vs *Xanthorrhoea*). Conversely, other pairs show high PR but low R² (e.g., *Epacris* vs *Grevillea* and *Banksia* vs *Epacris*), suggesting divergent spatial patterns of genetic structure and differing environmental drivers. These results suggest that similarity in putative environmental drivers is associated with similarity in spatial patterns of putative adaptive genomic variation across species within a shared landscape, although the strength of this relationship varies among species pairs.

**Figure 5.**
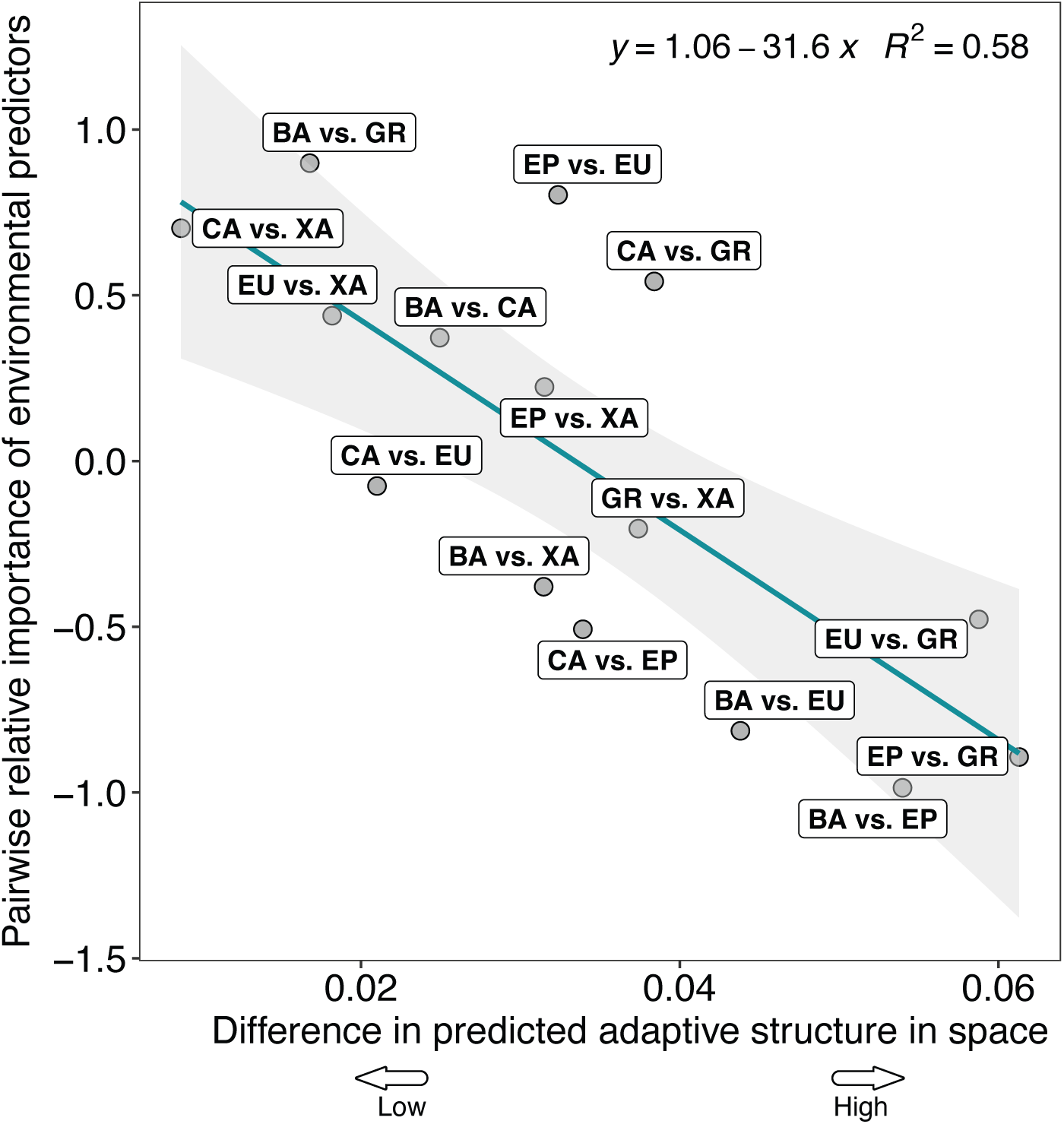
Scatter plot showing pairwise Pearson’s correlation between the mean relative importance (R2) of the predictor variables between species pairs and pairwise comparisons of predicted spatial genomic composition based on Procrustes residuals. Labels indicate the corresponding species pairs. The solid green line indicates the fitted regression slope. Labels indicate the corresponding species pairs. Species abbreviations are as follows: BA, *Banksia*; CA, *Callitris*; EP, *Epacris*; EU, *Eucalyptus*; GR, *Grevillea*; and XA, *Xanthorrhoea*.

## 4. Discussion

In a rare multi-species population-genomic analysis of the heterogeneous, mountainous landscape (Gariwerd in south-eastern Australia), we mapped the spatial patterns and environmental drivers of putatively adaptive genomic variation across co-occurring plant species with diverse life histories. Three key findings have emerged. First, we detected signals consistent with climate adaptation at short spatial scales, identifying putatively environmentally adapted loci across multiple, co-occurring, unrelated species with diverse life histories. Second, as expected, environmentally induced genomic responses varied among species, with some showing strong signals and others showing weaker or different patterns. Third, most species share overlapping high-elevation allelic turnover, which, combined with evidence of restricted gene flow across elevation gradients, may have implications for how species respond to environmental change. In aggregate, these results suggest that community-level genomic variation can be shaped by common environmental processes at fine spatial scales, while genomic responses can vary across species, reflecting a mixture of shared and species-specific patterns. We discuss the implications of these findings for biodiversity management under rapid environmental change.

### 4.1 Evidence of local adaptation across species at fine spatial scales

Fine-scale genomic signals of adaptation to climatic and soil variables were detected across all six unrelated plant species from Gariwerd, indicating that divergent selection operates over short spatial scales in this topographically complex landscape (Table S4). GEA analyses identified 1–4% of loci per species with significant environment-allele frequency relationships (Figures S3-S8), and GF analyses showed that 45–73% of these candidates had positive predictive power (Table S5). Collectively, these findings provide strong evidence that divergent selection is driving genomic differentiation across environmental gradients at microgeographic scales, consistent with other documented cases of local adaptation in plants and animals (Hargeby et al., 2004; Antonovics, 2006; Richardson & Urban, 2013; Richter-Boix et al., 2013; Yadav et al., 2021).

Although it is generally expected that climate- and soil-related variables collectively promote genetic divergence in plants (Hufford & Mazer, 2003; VanWallendael et al., 2019; Rupprecht et al., 2021), our results reveal that their relative importance varies markedly among species. Some species showed similar responses while others diverged in response to the same environmental gradients (Figure 5). Such differences have been observed in other systems, where closely related or co-occurring species vary in their signatures of local adaptation, reflecting differences in ecological traits, demography, genetic architecture, and other factors (Prates et al., 2018; Voolstra et al., 2023). Taken together, these patterns reflect species-specific interactions between environmental gradients and available genetic variation, even within shared landscapes (Goulet-Scott et al., 2024).

Across all species, the most pronounced genomic turnover of putatively adaptive alleles occurred in high-elevation habitats characterised by low temperature, low soil pH, low soil moisture, and high phosphorus levels (Figure 3; Table S6). Gradient forest identified soil-related variables as the most important predictors among the environmental variables tested in all species except *Eucalyptus* (Figure 2), suggesting that edaphic gradients associated with elevation contribute to adaptive genomic differentiation alongside climate. Soil properties are key drivers of plant community composition in Gariwerd (Enright et al., 1994; Pollock et al., 2015) and, therefore, are also likely to play an important role in adaptive evolutionary responses within species. These findings build upon previous studies, indicating that fine-scale environmental adaptation is common (Byars et al., 2007; Hoffmann et al., 2009; Anderson et al., 2015; Von Takach et al., 2021; Yadav et al., 2021; Gómez Quijano et al., 2024), despite the assumption that high gene flow often limits adaptive divergence in natural populations. Overall, the complex arrangement of fine-scale topographic features and environmental gradients across this region promotes adaptive divergence even across distances of hundreds to thousands of metres, a pattern consistent with selection exceeding the homogenising effects of gene flow in rugged terrain.

### 4.2 Variation in species responses to environmental features

Despite some similarities in putative environmental drivers across species, GF results indicate that species differ in the shape and steepness of their genetic responses to those drivers (Figures 2–4). Cumulative importance plots show that some species exhibit steep allelic turnover along particular gradients, whereas others display more gradual, near-linear shifts (Figure 2B). These patterns suggest that species respond to common selective gradients in different ways, yielding distinct spatial and environmental mosaics of allelic turnover (Figure 2). Such differences are expected given factors such as species-specific genetic architectures (e.g., effect-size distributions, pleiotropy, epistasis, linkage) (Yeaman & Whitlock, 2011; Whiting et al., 2024), demography and dispersal (influencing the balance between gene flow, drift and local selection) (Räsänen & Hendry, 2008; Pinho & Hey, 2010), life history and generation time (impacting the rate of evolutionary response) (Gandon & Michalakis, 2002; Raeymaekers et al., 2017), and ecological context (which can modulate the strength and form of selection) (Chaturvedi et al., 2022). Together, these factors shape changes in allele frequency along shared environmental gradients, influencing the form (e.g., gradual vs threshold-like), magnitude, and position of turnover. Broadly, comparisons across species pairs show that similarity in the importance of putative environmental drivers is associated with greater similarity in spatial patterns of genomic variation across the landscape. However, this relationship should be interpreted cautiously, as species were sampled across the same landscape and both metrics are derived from the same gradient forest models and a limited set of environmental variables, meaning that part of this pattern may reflect shared environmental structure and shared model structure.

### 4.3 Conservation implications

Most landscape genomic studies incorporating multiple species have largely focused on gene flow and population connectivity, showing that spatial patterns of neutral genetic structure are often conserved across taxonomically diverse taxa (Vandergast et al., 2008; Wood et al., 2013; Endo et al., 2015; Zbinden et al., 2023). These studies suggest that landscape features (e.g., geomorphology, habitat discontinuities, climatic gradients) often act as common barriers to gene flow despite differences in dispersal traits. In contrast, studies exploring how adaptive genomic variation relates to environmental factors across multiple species are still relatively limited (Hand et al., 2015; Koch et al., 2014; Zbinden et al., 2023). Our study indicates that while co-occurring species may respond to similar selective environments, the relative importance of environmental drivers can vary among taxa, producing both distinct and shared spatial patterns of putative adaptive variation across species pairs. These findings suggest that adaptive processes can be a primary driver of genomic variation over small spatial scales in complex landscapes, although adaptive genomic responses are not consistent across species. Consequently, predicting community-scale patterns of adaptive genomic variation, and potential vulnerabilities to environmental change, requires multi-species approaches that explicitly account for species-specific adaptive differences, rather than relying on a limited set of surrogate taxa.

Importantly, this study provides a framework for predicting maladaptation risk in Gariwerd plant communities. Genomic forecasts of maladaptation under future climates using genetic offset models have been criticised due to violations of key evolutionary assumptions and challenges in interpretation, particularly in the absence of supporting quantitative data (Rellstab et al., 2021; Lotterhos, 2024; Fitzpatrick et al., 2025; Ahrens et al., 2026). Instead, combining environmental genomic patterns with connectivity data may help identify species that differ in their capacity to respond to environmental change. Although gene flow can constrain local adaptation, it may also facilitate adaptation under environmental change by introducing pre-adapted genotypes (Sexton et al., 2014; Sacristán-Bajo et al., 2025). Here, we show that turnover in putatively adaptive alleles was greatest in high-elevation habitats, where populations are likely to be isolated and facing risks of maladaptation.

Previous work indicates that gene flow across elevation gradients in Gariwerd is limited for several focal species (Pollock et al., 2013; Lobos, 2024), suggesting that isolated high-elevation populations may be particularly vulnerable to climate change. In these systems, adaptation is likely to depend on standing genetic variation rather than the influx of pre-adapted alleles from non-local sources. Targeted interventions, such as assisted gene flow, may therefore be required to enhance the resilience of high elevation populations (Broadhurst et al., 2008; Aitken & Whitlock, 2013; Hoffmann et al., 2021). However, experimental validation (e.g., common garden trials) is required to confirm adaptive differentiation in climate-relevant traits and to verify the underlying selective drivers, thereby ensuring robust management recommendations.

### 4.4 Methodological considerations

Our multi-species genomic approach provides new insight into adaptive processes in complex landscapes, but methodological and conceptual limitations must be acknowledged. First, although reduced-representation sequencing offers an efficient and affordable strategy for generating genome-wide markers in non-model species, it samples only a fraction of loci across the genome, so many adaptive variants are likely to be missed (Lowry et al., 2017; da Fonseca et al., 2016; Capblancq et al., 2020). Second, species-specific genomic properties, such as variation in genome architecture, may explain differences in the ability to detect loci putatively under selection, even among taxa experiencing similar selection pressures (e.g., Hodgins & Yeaman, 2019). When multiple loci of small effect underlie a single polygenic trait, GEA approaches generally have low power to detect these associations (De Villemereuil et al., 2014). Third, although both BayPass and LFMM2 account for population structure, they can still yield spurious or missed associations when neutral demographic structure is aligned with adaptive divergence along environmental gradients, making it difficult to disentangle selection from history (Nadeau et al., 2016). Here, we analised candidate SNPs using GF to investigate the strength and direction of genomic shifts, and subsequently compared spatial and environmental turnover among species within a community analysis framework.

## 5. Conclusion

This study provides an in-depth multi-species genomic comparison of local adaptation across a topographically and climatically heterogeneous landscape. Despite sharing the same environmental mosaic, co-occurring plant species in Gariwerd showed both similar (high-elevation vulnerability) and species-specific genomic responses to shared environmental gradients, with similarities in environmental drivers broadly associated with spatial genomic patterns. Differences in the shape and magnitude of these responses highlight that common environmental gradients can generate diverse genomic outcomes across taxa. These results highlight that genomic responses to shared environmental gradients are not uniform across species, reinforcing the need to account for species-specific differences when applying genomic data in conservation contexts. Future work integrating genomic, experimental, and ecological data will help to clarify the mechanisms underlying these patterns and whether selection occurs predictably. Collectively, this knowledge would be key to determining the level of investment needed to assess resilience at community and ecosystem scales. Conservation of biodiversity under global change requires a community-level understanding of drivers of adaptive capacity.

## Author Contributions

A.D.M. received funding. A.D.M. and S.E.L. conceived and designed the study, with input and feedback from C.W.A. and P.D.R. A.D.M. and S.E.L. conducted the field work.

S.E.L. performed the bioinformatic and statistical analyses with significant input from C.W.A. S.E.L., K.A.H. and A.D.M. drafted the manuscript. All authors read, edited and approved the final manuscript.

## Acknowledgements

We thank Erlania, Emily Armstrong, Zach Clark, Harry Coleman, Jessica Fish, Owen Holland, and Daniel Vairo from Deakin University for their support during sample collections. A special thanks to Bill Weatherly (Friends of Forgotten Woodlands Inc.), Adam Merrick (Trust for Nature) and Simon Heyes (La Trobe University) for assistance with species and site selection.

## Ethical Statement

Permission to collect tissue samples was provided by DEECA under permit number 10010169 and adhered to all local guidelines.

## Benefit Sharing Statement

This research contributes to the conservation of flora in Gariwerd by openly sharing its findings with DEECA to inform and support local management efforts. Additionally, all data and results are publicly accessible, as detailed in the Data Availability Statement.

## Data Availability Statement

All data, along with the code required to reproduce the reported results are openly accessible via the Dryad Data Repository and the GitHub repository (https://github.com/selobos/Comparative-Environmental-Adaptation; https://zenodo.org/records/20845014). Raw sequence reads have been deposited in the National Centre for Biotechnology Information (NCBI) Sequence Read Archive (https://www.ncbi.nlm.nih.gov/sra/PRJNA1484297).

