## Supplemental File for "Topographic complexity shapes adaptive genomic variation in co-occurring plant species"

**Table of Contents:**

| **Table S1.** Summary of life history traits for the species examined in the study. | Page 2 |
| --- | --- |
| **Table S2.** Total number of individuals per species within each sampling location (n=30)  used for GEA testing. | Page 3-4 |
| **Table S3.** Total number of sample locations and SNPs retained for GEA analyses for each  plant species. | Page 5 |
| **Table S4.** Total number of candidate loci identified as having significant correlations with  environmental covariables by BayPass and LFMM2 for each species. | Page 6 |
| **Table S5.** The number of loci which had an *R*^2^ value greater than zero, and the maximum,  minimum, and mean *R*^2^ values from the Gradient Forest models run on candidate loci per  species. | Page 7 |
| **Table S6.** Principal component analysis (PCA) results for gradient forest modelling. | Page 8 |
| **Table S7.** The minimum (Min), first quartile (1st Qu), median, mean, third quartile (3rd Qu)  and maximum (Max) of pairwise Procrustes residuals. | Page 9 |
| **Figure S1.** Environmental data for genotype-environment association (GEA) analyses across  sampling locations within Gariwerd. | Page 10 |
| **Figure S2.** Value of the cross-entropy criterion as a function of the number of populations  For each species. | Page 11 |
| **Figure S3-S8.** Results of genotype-environment association (GEA) analyses show the  Strength of associations between individual loci and environmental variables identified by  BayPass and LFMM2. | Page 12-17 |

**Table S1.** Summary of life history traits for the species examined in the study.

| **Species** | **Common name** | **Family** | **Life-form** | **Pollen Vector** |
| --- | --- | --- | --- | --- |
| *Banksia marginata* | Silver Banksia | Proteaceae | Tree | Birds, insects, small mammals |
| *Callitris rhomboidea* | Oyster Bay Pine | Cupressaceae | Tree | Wind, Parrots and cockatoos |
| *Epacris impressa* | Common Heath | Ericaceae | Shrub | Insects |
| *Eucalyptus obliqua* | Messmate Stringybark | Myrtaceae | Tree | Bees |
| *Grevillea aquifolium* | Holly Grevillea | Proteaceae | Shrub | Birds and insects |
| *Xanthorrhoea australis* | Austral Grass-tree | Asphodelaceae | Grass-tree | Bees |

**Table S2.** Total number of individuals per species within each sampling location (n=30) used for GEA testing. Numbers after the slash indicate the subset of individuals selected for DArTseq sequencing. Underlined numbers indicate that a specimen was sequenced twice as a technical replicate. Species abbreviations as follows: BA, *Banksia marginata*; CA, *Callitris rhomboidea*; EP, *Epacris impressa*; EU, *Eucalyptus obliqua*; GR, *Grevillea aquifolium*; and XA, *Xanthorrhoea australis*.

| **Locations** | **Code** | **Species** | | | | | | **Elev.**  **(m)** | **Lat.**  **(ºS)** | **Lon.**  **(ºE)** |
| --- | --- | --- | --- | --- | --- | --- | --- | --- | --- | --- |
|  |  | **BA** | **CA** | **EP** | **EU** | **GR** | **XA** |  |  |  |
| Redman Rd | 1 | 4/6 | 4/6 |  | 1/6 | 6/6 | 6/7 | 344 | -37.2262 | 142.5511 |
| Yarram Gap Rd | 2 | 5/7 | 5/6 | 5/7 | 5/7 | 6/7 | 6/6 | 307 | -37.4550 | 142.5023 |
| Henham Track | 3 | 4/6 | 6/6 | 6/6 | 6/6 | 6/6 | 6/7 | 312 | -37.4080 | 142.4483 |
| Grampians Rd | 4 | 4/6 | 6/6 | 4/6 | 5/6 | 6/6 | 6/6 | 259 | -37.5214 | 142.4159 |
| Mackenzie Rd | 5A | 3/6 |  | 3/6 | 2/6 | 6/6 | 6/6 | 454 | -37.1178 | 142.4232 |
|  | 5C |  | 6/6 |  |  |  |  | 571 | -37.1360 | 142.4340 |
| Redman Rd | 6A | 6/6 | 6/7 | 6/7 |  |  | 6/6 | 431 | -37.2499 | 142.6076 |
|  | 6C |  |  |  |  | 6/6 |  | 452 | -37.2240 | 142.5850 |
| Mt Difficult Rd | 7A |  | 6/6 |  |  | 6/7 | 6/6 | 534 | -37.0419 | 142.4798 |
|  | 7B | 6/7 |  |  | 6/6 |  |  | 474 | -37.0364 | 142.4717 |
| Mt Zero Rd | 8A | 5/6 |  | 6/6 | 1/6 |  | 6/6 | 248 | -37.1111 | 142.5246 |
|  | 8B |  | 6/6 |  |  | 6/6 |  | 249 | -37.1050 | 142.5240 |
| Mt Zero Rd | 9 | 6/7 | 6/6 | 6/6 | 6/6 | 6/7 | 6/6 | 248 | -37.0559 | 142.5129 |
| Mt Zero Rd | 10 | 6/7 | 6/7 |  | 3/6 | 6/6 | 6/6 | 228 | -36.9159 | 142.4270 |
| Pohlners Rd | 11A |  | 5/6 |  |  | 6/6 | 6/6 | 389 | -36.9320 | 142.4047 |
|  | 11B |  |  |  | 5/6 |  |  | 365 | -36.9380 | 142.4080 |
|  | 11C | 6/6 |  | 5/6 |  |  |  | 369 | -36.9442 | 142.4109 |
| Pohlners Rd | 12 | 6/6 | 5/7 | 5/6 | 6/7 | 6/6 | 6/6 | 436 | -37.0135 | 142.4110 |
| Harrops Track | 13A | 5/6 | 6/6 | 6/6 |  | 6/6 | 4/7 | 236 | -37.2853 | 142.2618 |
|  | 13B |  |  |  | 6/6 |  |  | 239 | -37.2980 | 142.2570 |
| Harrops Track | 14A | 3/6 | 6/6 | 6/7 | 5/6 |  | 6/6 | 284 | -37.3580 | 142.2070 |
|  | 14B |  |  |  |  | 6/6 |  | 266 | -37.3680 | 142.1990 |
| Bullawin Rd | 15A | 6/7 | 6/7 | 6/6 | 6/7 | 6/7 | 6/7 | 279 | -37.4844 | 142.2689 |
| Serra Rd | 16 | 5/7 | 5/6 | 6/6 | 6/7 | 6/7 | 5/7 | 264 | -37.2857 | 142.4259 |
| Boroka Lookout | 17A | 5/6 | 5/6 | 5/7 | 4/6 | 6/6 | 6/6 | 852 | -37.1260 | 142.5005 |
| Henham Track | 18A |  | 6/7 |  |  |  |  | 424 | -37.3080 | 142.4870 |
|  | 18B | 5/6 |  | 5/6 |  | 6/6 | 6/6 | 350 | -37.3064 | 142.4851 |
| Serra Range Fireline | 19 | 4/6 | 6/6 | 6/6 | 5/6 | 6/7 | 6/7 | 289 | -37.5019 | 142.3644 |
| Lodge Rd | 20 | 4/6 | 5/7 | 6/6 | 6/6 | 6/6 | 6/6 | 229 | -37.1767 | 142.3180 |
| Asses Ear Rd | 21A | 4/7 | 6/6 |  |  | 6/6 | 6/6 | 228 | -37.1370 | 142.2665 |
|  | 21B |  |  |  | 4/7 |  |  | 226 | -37.1590 | 142.2780 |
|  | 21C |  |  | 5/6 |  |  |  | 218 | -37.1620 | 142.2790 |
| Glenelg River Rd | 22A | 5/6 |  | 6/6 | 4/6 | 6/6 |  | 277 | -37.3446 | 142.3376 |
|  | 22C |  |  |  |  |  | 6/7 | 259 | -37.3180 | 142.3667 |
| Mt Rosea | 23A |  | 6/6 | 3/6 |  |  |  | 846 | -37.1880 | 142.5010 |
|  | 23B |  |  |  |  | 6/7 |  | 846 | -37.1870 | 142.4980 |
|  | 23C | 4/6 |  |  |  |  |  | 777 | -37.1870 | 142.4934 |
|  | 23E |  |  |  | 6/6 |  | 6/6 | 685 | -37.1812 | 142.4855 |
| Mitchell Rd | 24A | 5/6 | 3/6 | 5/6 | 6/6 |  | 6/7 | 333 | -37.2938 | 142.6280 |
|  | 24B |  |  |  |  | 6/6 |  | 357 | -37.2730 | 142.6140 |
| Grampians Rd | 25 | 3/7 | 4/6 | 6/6 | 5/7 | 6/6 | 5/6 | 372 | -37.5918 | 142.3627 |
| Mt Abrupt | 26A | 5/6 |  | 4/6 |  |  | 6/6 | 816 | -37.5976 | 142.3533 |
|  | 26B |  | 4/6 |  |  |  |  | 641 | -37.5910 | 142.3570 |
| Victoria Valley Rd | 27A | 5/6 | 5/6 |  | 6/7 | 6/6 | 6/6 | 221 | -37.6053 | 142.3203 |
|  | 27B |  |  | 6/6 |  |  |  | 270 | -37.6220 | 142.3290 |
| Mt Thackeray | 28A |  |  | 6/6 | 6/6 |  |  | 966 | -37.2960 | 142.3370 |
|  | 28B |  |  |  |  | 6/7 | 6/6 | 856 | -37.3122 | 142.3293 |
|  | 28C |  | 6/7 |  |  |  |  | 786 | -37.2810 | 142.3360 |
|  | 28D | 5/7 |  |  |  |  |  | 814 | -37.2755 | 142.3421 |
| Jimmy Creek Rd | 29 | 6/6 |  | 6/6 | 5/6 | 6/6 | 6/6 | 335 | -37.3728 | 142.5131 |
| Mt William | 30A | 5/6 |  |  |  |  |  | 1161 | -37.2927 | 142.6011 |
|  | 30B |  |  | 4/6 |  |  |  | 1076 | -37.2920 | 142.5970 |
|  | 30D |  |  |  |  |  | 6/6 | 1009 | -37.2909 | 142.5930 |
|  | 30E |  | 6/7 |  |  | 6/6 |  | 943 | -37.2870 | 142.5940 |
|  | 30F |  |  |  | 6/6 |  |  | 858 | -37.2830 | 142.5870 |

**Table S3.** Total number of sample locations and SNPs retained for GEA analyses for each plant species.

| **Species** | **Pre-filtering** | |  | **Post-filtering** | | |
| --- | --- | --- | --- | --- | --- | --- |
|  | **No. SNPs** | **No.**  **individuals** |  | **No. SNPs** | **No.**  **individuals** | **No. populations** |
| *Banksia marginata* | 20,212 | 188 |  | 2,521 | 145 | 30 |
| *Callitris rhomboidea* | 72,381 | 176 |  | 16,361 | 152 | 28 |
| *Epacris impressa* | 135,235 | 164 |  | 13,725 | 143 | 27 |
| *Eucalyptus obliqua* | 80,226 | 188 |  | 9,423 | 132 | 27 |
| *Grevillea aquifolium* | 190,026 | 182 |  | 32,834 | 174 | 29 |
| *Xanthorrhoea australis* | 140,212 | 188 |  | 35,082 | 174 | 30 |

**Table S4.** Total number of candidate loci identified as having significant correlations with environmental covariables by BayPass and LFMM2 for each species (including number of shared loci between analytical methods). *TWI*–topographic wetness index, *S*_AP_–available phosphorus, *S*_pH_–soil pH in water and CaCl_2_ and *T*_MAX_–maximum temperature of warmest month.

| **Species** | **Predictors** | **BayPass** |  | **LFMM2** |  | **Shared** |  | **Total** |  | **Unique** |
| --- | --- | --- | --- | --- | --- | --- | --- | --- | --- | --- |
| *Banksia marginata* | *TWI* | 6 |  | 1 |  | 0 |  | 7 |  | 7 |
|  | *S*_AP_ | 7 |  | 1 |  | 0 |  | 8 |  | 8 |
|  | *S*_pH_ | 7 |  | 4 |  | 1 |  | 11 |  | 10 |
|  | *T*_MAX_ | 0 |  | 5 |  | 0 |  | 5 |  | 5 |
| *Callitris rhomboidea* | *TWI* | 32 |  | 28 |  | 10 |  | 60 |  | 50 |
|  | *S*_AP_ | 34 |  | 19 |  | 5 |  | 53 |  | 48 |
|  | *S*_pH_ | 14 |  | 17 |  | 5 |  | 31 |  | 26 |
|  | *T*_MAX_ | 0 |  | 34 |  | 0 |  | 34 |  | 34 |
| *Epacris impressa* | *TWI* | 23 |  | 42 |  | 5 |  | 65 |  | 60 |
|  | *S*_AP_ | 21 |  | 24 |  | 2 |  | 45 |  | 43 |
|  | *S*_pH_ | 29 |  | 73 |  | 10 |  | 102 |  | 92 |
|  | *T*_MAX_ | 11 |  | 80 |  | 5 |  | 91 |  | 86 |
| *Eucalyptus obliqua* | *TWI* | 11 |  | 12 |  | 5 |  | 23 |  | 18 |
|  | *S*_AP_ | 22 |  | 9 |  | 6 |  | 31 |  | 25 |
|  | *S*_pH_ | 10 |  | 16 |  | 2 |  | 26 |  | 24 |
|  | *T*_MAX_ | 3 |  | 47 |  | 1 |  | 50 |  | 49 |
| *Grevillea aquifolium* | *TWI* | 83 |  | 108 |  | 29 |  | 191 |  | 162 |
|  | *S*_AP_ | 84 |  | 165 |  | 19 |  | 249 |  | 230 |
|  | *S*_pH_ | 65 |  | 43 |  | 12 |  | 108 |  | 96 |
|  | *T*_MAX_ | 17 |  | 100 |  | 5 |  | 117 |  | 112 |
| *Xanthorrhoea australis* | *TWI* | 19 |  | 32 |  | 3 |  | 51 |  | 48 |
|  | *S*_AP_ | 26 |  | 42 |  | 9 |  | 68 |  | 59 |
|  | *S*_pH_ | 18 |  | 55 |  | 9 |  | 73 |  | 64 |
|  | *T*_MAX_ | 17 |  | 49 |  | 6 |  | 66 |  | 60 |

**Table S5.** The number of loci which had an *R*^2^ value greater than zero, and the maximum, minimum, and mean *R*^2^ values from the Gradient Forest models run on candidate loci per species.

| **Species** | **No. of retained SNPs with *R*^2^ > 0** | **Max. *R*^2^ of retained SNPs** | **Min. *R*^2^ of retained SNPs** | **Mean *R*^2^ of retained SNPs** |
| --- | --- | --- | --- | --- |
| *Banksia marginata* | 17 | 0.153 | 2.25e^-04^ | 0.038 |
| *Callitris rhomboidea* | 80 | 0.271 | 1.49e^-04^ | 0.051 |
| *Epacris impressa* | 156 | 0.444 | 8.69e^-06^ | 0.066 |
| *Eucalyptus obliqua* | 65 | 0.302 | 2.79e^-04^ | 0.063 |
| *Grevillea aquifolium* | 250 | 0.357 | 8.84e^-05^ | 0.049 |
| *Xanthorrhoea australis* | 131 | 0.371 | 8.55^-05^ | 0.053 |

**Table S6.** Principal component analysis (PCA) results for gradient forest modelling. For each variable, the numbers in the row represent the strength of its correlation with the eigenvector of each principal component (PC). % Variance indicates the variance accounted for by each component of the total variance in all the environmental variables. Environmental variables ordered by their overall importance for predicting compositional turnover in allele frequency. Abbreviations of the variables are *TWI*–topographic wetness index; *S*_AP_–available phosphorus; *S*_pH_–soil pH in water and CaCl_2,_ and *T*_MAX_–maximum temperature of the warmest month.

| *Banksia marginata* | |  |  | *Eucalyptus obliqua* | |  |  |
| --- | --- | --- | --- | --- | --- | --- | --- |
|  | PC1 | PC2 | PC3 |  | PC1 | PC2 | PC3 |
| *S*_AP_ | **0.8825** | 0.4335 | -0.1310 | *TWI* | **-0.6764** | -0.5592 | -0.4793 |
| *T*_MAX_ | -0.3400 | **0.5289** | 0.1917 | *S*_pH_ | -0.4987 | **0.7350** | -0.1514 |
| *TWI* | -0.2227 | 0.1514 | **-0.9623** | *T*_MAX_ | -0.3194 | **0.3220** | 0.0701 |
| *S*_pH_ | -0.2368 | **0.7137** | 0.1417 | *S*_AP_ | 0.4379 | 0.2082 | **-0.8616** |
| % Variance | *65* | *19* | *11* | % Variance | *57* | *23* | *16* |
| *Callitris rhomboidea* | |  |  | *Grevillea aquifolium* | |  |  |
|  | PC1 | PC2 | PC3 |  | PC1 | PC2 | PC3 |
| *TWI* | 0.5957 | **-0.7598** | 0.2595 | *S*_AP_ | **0.8420** | -0.5039 | -0.1688 |
| *S*_AP_ | -0.5844 | -0.1891 | **0.7758** | *TWI* | -0.2411 | -0.0858 | **-0.9661** |
| *T*_MAX_ | 0.4079 | 0**.4343** | 0.2727 | *S*_pH_ | -0.3743 | **-0.7738** | 0.1796 |
| *S*_pH_ | 0.3705 | 0.4453 | **0.5063** | *T*_MAX_ | -0.3046 | **-0.3742** | 0.0773 |
| % Variance | *55* | *21* | *19* | % Variance | *59* | *27* | *9* |
| *Epacris impressa* | |  |  | *Xanthorrhoea australis* | |  |  |
|  | PC1 | PC2 | PC3 |  | PC1 | PC2 | PC3 |
| *S*_pH_ | **-0.6880** | 0.3989 | -0.5426 | *TWI* | 0.6134 | **-0.6145** | -0.4960 |
| *TWI* | -0.4040 | **-0.8729** | -0.2142 | *T*_MAX_ | 0.3822 | **0.4118** | -0.0253 |
| *T*_MAX_ | -0.5803 | 0.1926 | **0.6584** | *S*_AP_ | -0.5449 | 0.1145 | **-0.8131** |
| *S*_AP_ | 0.1635 | 0.2047 | **-0.4757** | *S*_pH_ | 0.4252 | **0.6631** | -0.3036 |
| % Variance | *63* | *23* | *9* | % Variance | *57* | *21* | *18* |

*Note:* In bold are the top significant loadings indicating how much of its variation is being summarised by the particular component.

**Table S7.** The minimum (Min), first quartile (1^st^ Qu), median, mean, third quartile (3^rd^ Qu) and maximum (Max) of pairwise Procrustes residuals. Species abbreviations as follows: BA, *Banksia marginata*; CA, *Callitris rhomboidea*; EP, *Epacris impressa*; EU, *Eucalyptus obliqua*; GR, *Grevillea aquifolium*; and XA, *Xanthorrhoea australis*.

|  | **Min** | **1^st^ Qu** | **Median** | **Mean** | **3^rd^ Qu** | **Max** |
| --- | --- | --- | --- | --- | --- | --- |
| CA. vs. XA | 0.00006 | 0.00159 | 0.00211 | 0.00233 | 0.00293 | 0.00871 |
| BA. vs. GR | 0.00012 | 0.00371 | 0.00508 | 0.00527 | 0.00653 | 0.01679 |
| EU. vs. XA | 0.00019 | 0.00601 | 0.00776 | 0.00797 | 0.00949 | 0.01819 |
| CA. vs. EU | 0.00011 | 0.00403 | 0.00503 | 0.0055 | 0.00651 | 0.02101 |
| BA. vs. CA | 0.00012 | 0.00367 | 0.00482 | 0.00511 | 0.00633 | 0.02494 |
| BA. vs. XA | 0.00015 | 0.00365 | 0.00503 | 0.00528 | 0.00652 | 0.03146 |
| EP. vs. XA | 0.00017 | 0.00614 | 0.00827 | 0.00914 | 0.01147 | 0.03151 |
| EP. vs. EU | 0.00022 | 0.00467 | 0.00668 | 0.00749 | 0.0097 | 0.03237 |
| CA. vs. EP | 0.00011 | 0.00439 | 0.00564 | 0.00656 | 0.00764 | 0.03392 |
| GR. vs. XA | 0.00008 | 0.00261 | 0.00396 | 0.00485 | 0.00562 | 0.03742 |
| CA. vs. GR | 0.00006 | 0.00265 | 0.00347 | 0.00446 | 0.00463 | 0.03839 |
| BA. vs. EU | 0.00023 | 0.00585 | 0.00777 | 0.008 | 0.00964 | 0.04381 |
| BA. vs. EP | 0.00054 | 0.00556 | 0.00805 | 0.00857 | 0.01075 | 0.05398 |
| EU. vs. GR | 0.00024 | 0.00577 | 0.00874 | 0.01052 | 0.01456 | 0.05876 |
| EP. vs. GR | 0.00113 | 0.00522 | 0.00818 | 0.01026 | 0.01402 | 0.02190 |

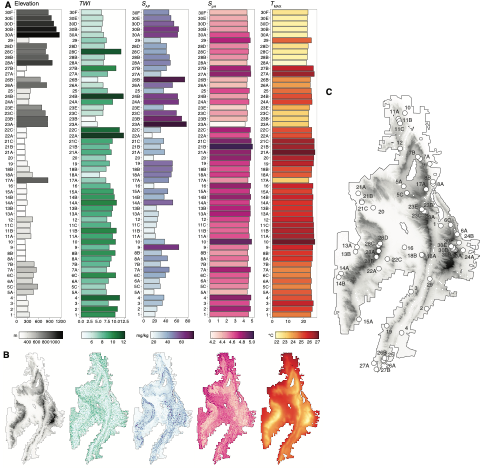
**Figure S1.** Environmental data for genotype-environment association (GEA) analyses across sampling locations within Gariwerd. (A) Bar plots representing the estimates of each variable across the 30 unique sampling locations. (B) Map illustrating the spatial distribution of each variable across Gariwerd. (C) Map displaying the 30 sampling locations, each indicated by a small white circle (see Table 3.S2 for location names).

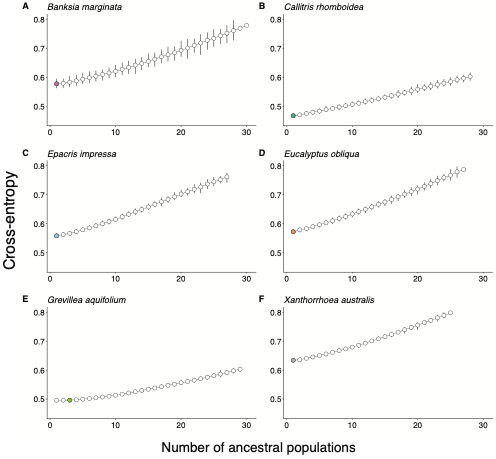
**Figure S2.** Value of the cross-entropy criterion as a function of the number of populations for each species.

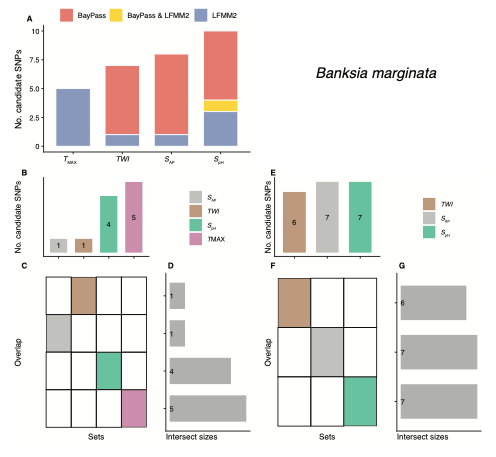
**Figure S3.** Results of genotype-environment association (GEA) analyses show the strength of associations between individual loci and environmental variables identified by BayPass and LFMM2. (A) Bar plot shows the number of putative candidate loci associated with each environmental variable. UpSet plots of the candidate loci found in LFFM2 (left; B–D) and BayPass (right; E–G), respectively. (B) & (E) Bar plots show the total number of candidate loci found in each environmental variable. (C) & (F) Heatmaps below the bar plot indicate which sets of candidate loci are represented by each bar. (D) & (G) Stacked bar plots show the total number of individual candidate loci identified in one or more environmental variables.

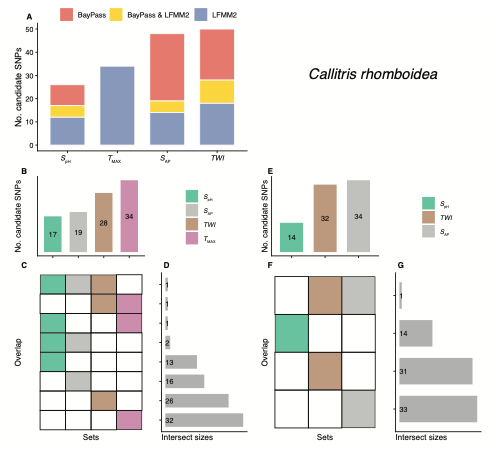
**Figure S4**. Results of genotype-environment association (GEA) analyses show the strength of associations between individual loci and environmental variables identified by BayPass and LFMM2. (A) Bar plot shows the number of putative candidate loci associated with each environmental variable. UpSet plots of the candidate loci found in LFFM2 (left; B–D) and BayPass (right; E–G), respectively. (B) & (E) Bar plots show the total number of candidate loci found in each environmental variable. (C) & (F) Heatmaps below the bar plot indicate which sets of candidate loci are represented by each bar. (D) & (G) Stacked bar plots show the total number of individual candidate loci identified in one or more environmental variables.

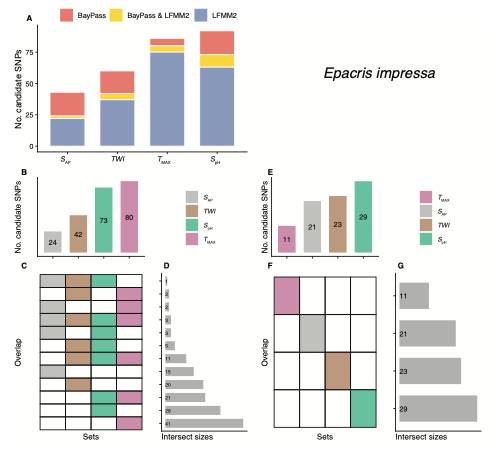
**Figure S5.** Results of genotype-environment association (GEA) analyses show the strength of associations between individual loci and environmental variables identified by BayPass and LFMM2. (A) Bar plot shows the number of putative candidate loci associated with each environmental variable. UpSet plots of the candidate loci found in LFFM2 (left; B–D) and BayPass (right; E–G), respectively. (B) & (E) Bar plots show the total number of candidate loci found in each environmental variable. (C) & (F) Heatmaps below the bar plot indicate which sets of candidate loci are represented by each bar. (D) & (G) Stacked bar plots show the total number of individual candidate loci identified in one or more environmental variables.

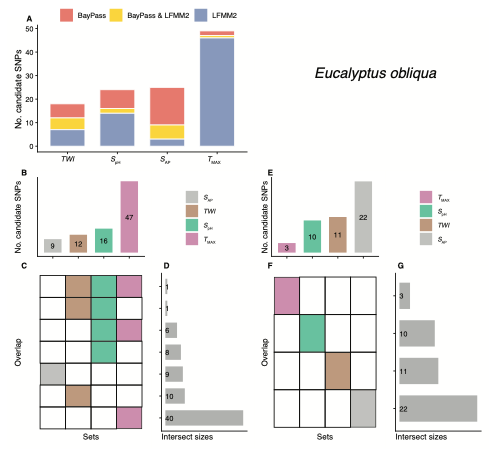
**Figure S6.** Results of genotype-environment association (GEA) analyses show the strength of associations between individual loci and environmental variables identified by BayPass and LFMM2. (A) Bar plot shows the number of putative candidate loci associated with each environmental variable. UpSet plots of the candidate loci found in LFFM2 (left; B–D) and BayPass (right; E–G), respectively. (B) & (E) Bar plots show the total number of candidate loci found in each environmental variable. (C) & (F) Heatmaps below the bar plot indicate which sets of candidate loci are represented by each bar. (D) & (G) Stacked bar plots show the total number of individual candidate loci identified in one or more environmental variables.

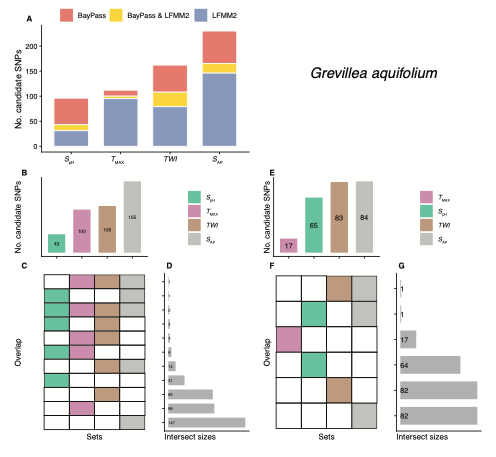
**Figure S7.** Results of genotype-environment association (GEA) analyses show the strength of associations between individual loci and environmental variables identified by BayPass and LFMM2. (A) Bar plot shows the number of putative candidate loci associated with each environmental variable. UpSet plots of the candidate loci found in LFFM2 (left; B–D) and BayPass (right; E–G), respectively. (B) & (E) Bar plots show the total number of candidate loci found in each environmental variable. (C) & (F) Heatmaps below the bar plot indicate which sets of candidate loci are represented by each bar. (D) & (G) Stacked bar plots show the total number of individual candidate loci identified in one or more environmental variables.

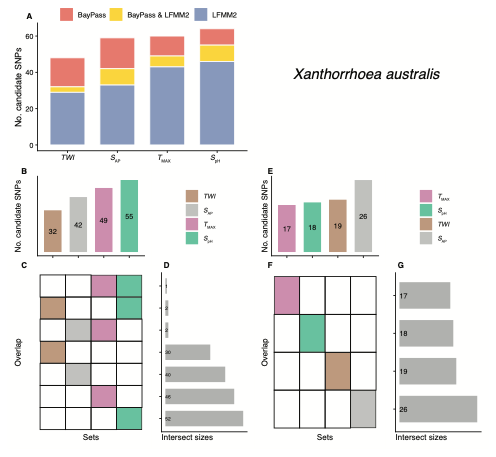
**Figure S8.** Results of genotype-environment association (GEA) analyses show the strength of associations between individual loci and environmental variables identified by BayPass and LFMM2. (A) Bar plot shows the number of putative candidate loci associated with each environmental variable. UpSet plots of the candidate loci found in LFFM2 (left; B–D) and BayPass (right; E–G), respectively. (B) & (E) Bar plots show the total number of candidate loci found in each environmental variable. (C) & (F) Heatmaps below the bar plot indicate which sets of candidate loci are represented by each bar. (D) & (G) Stacked bar plots show the total number of individual candidate loci identified in one or more environmental variables.
